# The Cxcl14 chemokine defines pioneer axon guidance and early circuit assembly in the inner ear

**DOI:** 10.64898/2026.05.21.726833

**Authors:** M Rumbo, A Bañón, A Nechiporuk, B Alsina

## Abstract

The nervous system wiring requires the precise coordination of axon guidance, neuronal migration, and target cell recognition. Here, we show that inner ear circuit formation, relies on pioneer cells extending an axonal scaffold that selectively target nascent hair cells. High spatiotemporal imaging of pioneer axons reveal how they navigate through the cranial environment, establish dynamic cell-cell contacts with other axons to finally stabilize in target cells. These pioneer axons are required not only for follower axon growth but also for coordinated migration of follower neurons, revealing a cellular hierarchy underlying circuit assembly. We identify the chemokine Cxcl14 as a novel instructive guidance cue regulating pioneer axon extension, turning, and fasciculation at discrete cellular decision points. Loss of Cxcl14 disrupts axonal navigation, compromises synaptic organization in hair cells, and impairs mechanosensory-based behavior. Together, our findings establish a new chemokine-based mechanism linking pioneer axon guidance to early sensory circuit assembly necessary for building mechanosensory networks.

## Introduction

The establishment of precise neural circuits during embryogenesis requires a coordinated sequence of neuronal migration, axon extension and navigation through complex molecular environments, combined with selective target recognition to ensure accurate signal transmission. A highly conserved mechanism for circuit development is the extension, first, of pioneer axons that organize a first scaffold circuit that is used for migration of later born neurons or growth of follower axons (Hidalgo and Brand, 1997; Supèr *et al*., 1998).

The inner ear is a key sensory organ of our head responsible for detecting and transmitting acoustic, linear and rotational acceleration information to the brain. This is achieved through the establishment of a mechanosensory circuit connecting hair cells (HCs) with bipolar neurons of the statoacoustic ganglion (SAG). HCs are strategically arranged in separated sensory domains within the otic epithelium to detect sensory information across the three orthogonal head axes (Fritzsch *et al*., 2002; Higuchi *et al*., 2019). As SAG neurons are placed outside the otic epithelium, axons must navigate within the cranial mesenchyme and find their target HCs. Several works have addressed the molecular mechanisms regulating SAG axonal guidance towards HCs. The generation of ectopic HCs in the inner ear epithelium of mice embryos resulted in rerouting of SAG axons and targeting of the ectopic HCs, suggesting the secretion of a putative attractive molecule by HCs (Zheng and Gao, 2000). Although BDNF and NT3 neurotrophins, expressed in all HCs, were postulated to be the attractant signaling molecules, it was difficult to decipher whether they exerted trophic or tropic roles on SAG axons (Tessarollo, Coppola and Fritzsch, 2004). Conversely, in chick otic explants, SAG axons reached sensory regions before HC differentiation (Hemond and Morest, 1992). Similarly, in mouse mutants lacking BDNF and HC differentiation, SAG axons still reached the sensory domains, suggesting that other earlier-expressed molecules in the sensory region could mediate the early guidance of SAG axons (Fritzsch *et al*., 2005). Taking these results into consideration, the guidance mechanisms by which SAG axons reach early-born HCs located in distinct regions are yet poorly understood. In addition, whether the establishment of this circuitry relies in pioneer axons remains to be explored.

Recently, we identified a population of non-otic pioneer SAG neurons (thereafter, pioneers) that are necessary for the migration and coalescence of otic neuroblasts into distinct lobes (Bañón and Alsina, 2023). While pioneers of cranial ganglia have received less attention, studies on the posterior lateral line (LL), a mechanosensory system that senses water flow through HCs organized in neuromasts and develops in close contact to the otic vesicle, demonstrated that pioneer axons targeted the migrating primordium, guiding follower axons (Ghysen and Dambly-Chaudière, 2004). These posterior LL (pLL) pioneers were characterized by a specific transcriptomic signature and the usage of *ret-gfra1* receptors to mediate axonal growth and dynamics (Sato and Takeda, 2013; Tuttle *et al*., 2019; Woodruff *et al*., 2025).

Chemokines are mainly described to direct immune cell migration (Hughes and Nibbs, 2018), but recent studies have explored other roles in cell differentiation, survival, synaptic activity and axon guidance (Li *et al*., 2005; Borrell and Marín, 2006; Rostène *et al*., 2011). The Cxcl12-Cxcr4 signalling axis, for instance, is present in organisms before the appearance of an immune system and is a regulator of neural development (Li and Ransohoff, 2008). Cxcl14 is one of the closest homologs of Cxcl12, implicated mainly in angiogenesis, tumor growth and migration of immune cells in adults. However, its functions remain comparatively unexplored, largely because it does not seem to interact with the classical CXC-type receptors and its cognate receptor has not yet been identified (Tanegashima *et al*., 2013; Collins *et al*., 2017; Al Hamwi *et al*., 2024). In the mouse hippocampus, *Cxcl14* was described to regulate the differentiation of rare microglia (Schmid *et al*., 2009; Li, Li and Jiao, 2019). Moreover, Cxcl14 has been implicated in the regulation of GABAergic synaptic transmission and differentiation of cortical interneurons, modulating neuronal activity and circuit maturation (Banisadr *et al*., 2011; Iannone *et al*., 2024). Intriguingly, Cxcl14 shows a highly conserved expression pattern in HCs of the inner ear in zebrafish, *Xenopus*, mouse and human (Park *et al*., 2009; Gordon *et al*., 2011; Wang *et al*., 2024), and it was also found to be enriched in regenerating HCs after damage in chick and mouse, assessed by scRNA-seq (Janesick *et al*., 2022; Luca *et al*., 2025). Its evolutionary conserved expression in HCs and its role in synaptic activity and circuit maturation, suggests a putative role in inner ear circuit formation.

In this work, we reveal that SAG pioneers share some transcriptional markers with anterior LL (aLL) progenitors, suggesting a common SAG-aLL pioneer pool involved in the development of both cranial ganglia. We also provide for the first time high spatiotemporal imaging of pioneer axonal dynamics, showing their first contact with HCs and their requirement for the migration of later migrating otic neuroblasts. Overall, these results highlight a fundamental role of pioneer axons is the establishment of the scaffold for the organization of the future mechanosensory circuit. Finally, when investigating the molecular cues for this process, we identified the Cxcl14 chemokine as a central cue regulating pioneer axon guidance. In *cxcl14*-deficient embryos, SAG pioneer axons exhibited pathfinding errors, reduced fasciculation and turning errors. At later developmental stages, branching complexity at innervation sites was diminished, accompanied by incorrect HC targeting, a reduction of HC activity and synaptic disorganization. Moreover, *cxcl14*-deficient embryos also displayed strong phenotypes in the LL and SAG central fascicles projecting to the hindbrain. Disruption of the circuitry, resulted in a significantly reduced startle response, indicating a functional impairment of the vestibuloacoustic system.

Together, our findings reveal the existence and relevance of pioneer axonal circuit assembly in the inner ear and we identify Cxcl14 as a unique molecule regulating both peripheral and central projections of SAG pioneer axons. This chemokine emerges as a potential candidate to mediate reinnervation of HCs after injury.

## Results

### Pioneer SAG neurons share a transcriptional profile with aLL progenitors

We recently identified a group of non-otic pioneer cells, positive for *neurod1*, positioned anterior to the otic primordium that orchestrate the migration and coalescence of later inner ear delaminating neuroblasts (Bañón and Alsina, 2023) (Fig 1A). However, the transcriptomic identity of these pioneer cells has not been addressed. As SAG pioneers develop next to the anterior LL ganglion (aLL), we wondered whether the SAG pioneers and aLL progenitors could share their lineage and transcriptomic profile. To assess if this is the case, we analyzed scRNA-seq data sets from the LL published in Woodruff *et al*., 2025 to identify aLL neurons and their progenitors from embryos at distinct stages. We subclustered previously annotated LL cells and used differential expression analysis and previously defined markers to identify aLL and pLL neurons as well as their progenitor populations (Fig. 1B, C). For example, pLL lineage displays high levels of *hoxb5b*, whereas aLL progenitors and aLL neurons express high levels of *alcama* compared to the pLL (Fig. 1C). As SAG pioneers are evident at 18 hpf, we asked whether any known SAG genes are expressed in the developing aLL at this stage, potentially marking SAG pioneers. Differential expression analysis identified several aLL transcripts including *pou2f2a.1*, *irx1a* and *irx1b*, *greb1l*, *itga5*, and *fstl1a* (p adjusted value < 0.001; Fig. 1D, E). We assessed whether any of these markers were expressed in SAG pioneers by in situ hybridization, which confirmed *irx1a* expression in SAG pioneers but not all the aLL progenitors (Fig. 1F and F’). Moreover, we found Ret-Gfra1 receptors expression highly present in the SAG pioneers (Fig. 1G). Altogether, we find that SAG pioneers share markers with other sensory pioneers, suggesting that pioneer identity represents a conserved cellular program deployed across distinct mechanosensory systems.

**Fig 1.**
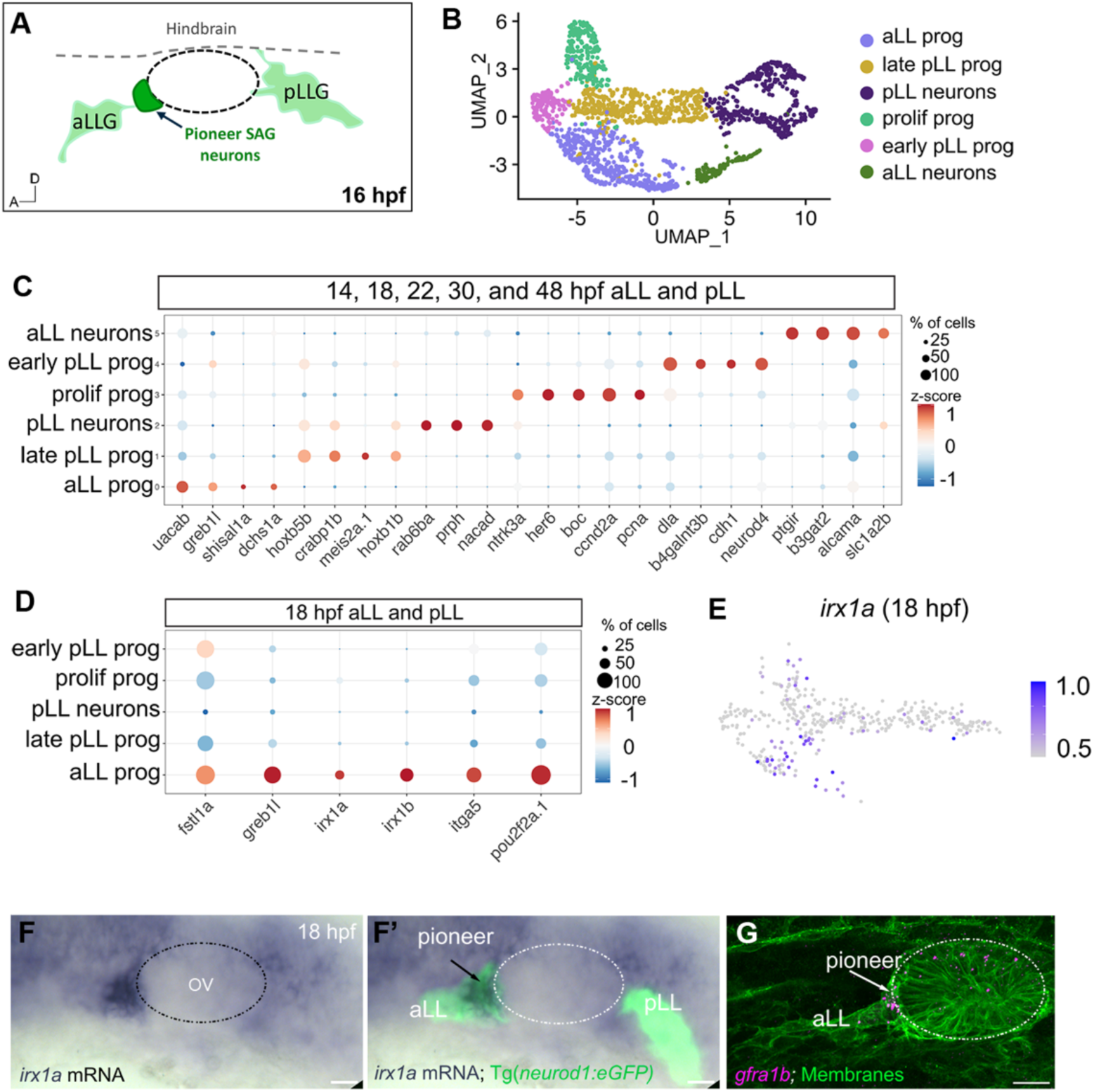
SAG pioneers share a transcriptional profile with aLL progenitors. (A) Schematic representation of pioneer SAG neurons (dark green), aLL and pLL ganglia (light green) at 16 hpf. Photoconversion from green to magenta of the SAG pioneers using the Tg(*neurod1:kikume*) line. Visualization of the photoconverted neurons at 24 hpf, in which SAG pioneer neurons (magenta) extend the first pioneer axons of the SAG, targeting the posterior region of the OV (white arrow labelling growth cone). (B) UMAP of lateral line, TgBAC(*neurod1:eGFP*)^nl1^-positive cells from 14, 18, 22, and 48 hpf clustered as aLL progenitors, late pLL progenitos, pLL neurons, proliferating progenitors, early pLL progenitors and aLL neurons. (C) Dotplot showing expression of enriched markers in each subcluster. (D) Dotplot showing expression of known SAG markers in the lateral line in the 18 hpf data set. (E) Feature plots showing expression of irx1a at 18 hpf. (F-F’) *irx1a* expression in pioneers (dark blue) with sensory neurons visualized by TgBAC(*neurod1:eGFP*)^nl1^ (green) at 18 hpf. (G) FISH labelling *gfra1b* (magenta), a pioneer marker. Membranes are labelled by *cldnB:memgfp* transgene (green). aLLG: anterior lateral line ganglion; pLLG: posterior lateral line ganglion, SAG: statoacoustic ganglion; OV: otic vesicle. Dotted circle represents OV.

### Pioneer neurons extend pioneer axons that target first born HCs and serve as scaffolds for the migration of otic neuroblasts

We then decided to explore whether the pioneer SAG neurons establish the first scaffolding axonal circuit. For this, we labeled the pioneer cells via photoconversion. Indeed, the first axons extending posteriorly were exclusively marked by a photoconverted fluorophore, confirming their pioneer identity (Fig. 2A, white arrow). To better understand how pioneer axons extend to HCs, we performed confocal time-lapse acquisitions at high spatiotemporal resolution. We focused on the extension of the pioneer SAG axons to the posterior region of the otic vesicle (OV). Initially, one or two neurites extended posteriorly, arrowhead marks the axon growth cone; Fig. 2B, Video 1). During this period, the extending pioneer axon also produced many shorter perpendicular branches that finally retracted (Fig. 2C, red arrows; Video 2). We confirmed that these pioneer axons contacted the first-born HC by photoconverting the SAG pioneers using the Tg(*neurod1:kikume*) line and simultaneously labelling HC with the Tg(*Brn3C:GFP*) *^s356t^*reporter (Fig. 2D). A thickening of the axon near the posterior newborn HC was visible, indicative of the first axon-HC contact (Fig. 2C and D, yellow circles). Surprisingly, the SAG pioneer axon did not stop in that thickened contact but extended posteriorly, passing past the inner ear and establishing thinner contacts with the hindbrain and axonal branches from the pLL central projection (Fig. 2C and D, green arrowheads). The pioneer SAG axon-pLL axon interaction finally resolved (Fig Suppl. 1, Video Suppl. 1). The process of SAG pioneer axon extension, branching/refinement, retraction and final stabilization to the HC lasted 5 hours, from 20 to 25 hpf.

**Fig. 2.**
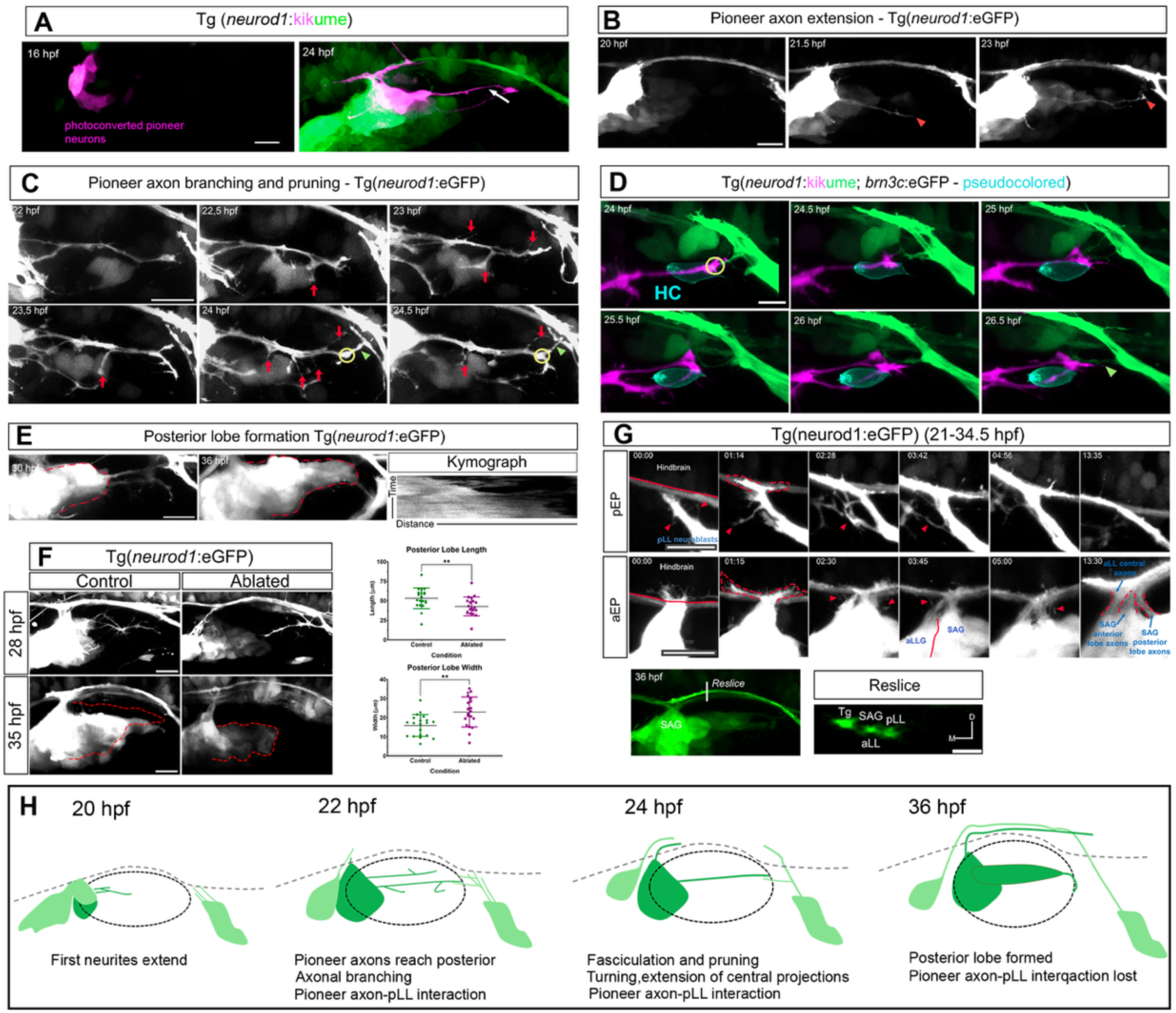
SAG pioneer axons target newborn HC and serve as scaffolds for neuroblast migration. Central projections grow simultaneously in a stereotyped manner. (A) Photoconversion from green to magenta of the SAG pioneers using the Tg(*neurod1:kikume*) line. Visualization of the photoconverted neurons at 24 hpf, in which SAG pioneer neurons (magenta) extend the first pioneer axons of the SAG, targeting the posterior region of the OV (white arrow labelling growth cone). (B) Representative stills from a confocal time-lapse of the SAG pioneer axon growth to the posterior region of the OV from 20 to ≈24 hpf. Sensory neurons and axons are visualized by TgBAC(*neurod1:*eGFP)^nl1^. Scale bar = 20 µm. Red arrowheads mark the growth cone of the axon. (C) Representative stills of a confocal time-lapse showing branching and pruning of SAG pioneer axons (red arrows) from 22 to 24.5 hpf. Yellow circle indicates the thickening of the axon when the posterior sensory region is reached. Green arrowhead indicates the contact between SAG pioneer axons and pLL axons. Scale bar = 10 µm (D) Photoconverted SAG pioneer axon (magenta) targets and contacts the first newborn HC (pseudocolored in cyan). A thickening in the axon is observed at the base of the HC (yellow circle). pLL (green) axons that contact SAG pioneer axons are depicted by a green arrowhead. Scale bar = 10 µm. (E) Stills showing the crawling of neuroblasts along the SAG pioneer axons to form the posterior lobe (marked with a red dotted line). Kymograph representing how the axon gets populated by neuroblasts through time (F) Confocal images from control and pioneer neuron-ablated embryos at 28 and 35 hpf. Arrow indicates pLL axons entering otic territory. Red dotted line indicates posterior lobe shape. Quantifications for posterior lobe length and width are shown in the graphs on the rigth. N = 20 controls 20 ablated. Lobe length: control = 53.18 ± 13.2 and ablated = 42.72 ± 12.2, **p-value = 0.0039. Lobe width: control = 15.85 ± 5.7 and ablated = 22.98 ± 7.8, **p-value = 0.0026. Scale bar = 20 µm. (G) Still confocal images from a time series showing the growth and entrance to the hindbrain of central projections from pLL and aLL-SAG, respectively, in the hindbrain entry points between 22 and 35.5 hpf. Arrowheads = temporal branches. Dashed lines are used to mark central axons when they turn. Scale bar = 20 µm. Confocal image of a 36 hpf SAG from a lateral view and the orthogonal view (reslice) of the fasciculated central projections of Tg, aLL, SAG and pLL. (H) Schematic representation of the dynamics of growth of the SAG and the central projections from aLL and pLL from 20 to 36 hpf. pEP: posterior Entry Points; aEP: anterior Entry Points.

Remarkably, starting at 24 hpf, newly delaminated neuroblasts crawled along the pioneer axons to form the posterior lobe (Fig. 2E, posterior lobe marked with a red dotted line; Video 3). To address whether pioneer axons were required for this process, we ablated SAG pioneers at 16 hpf, which resulted in the absence of the pioneer axons at 28 hpf. In some ablated embryos, we observed a long branch of the pLL central bundle extending into the otic domain (Fig. 2F, red arrowhead). Quantification of the posterior lobe revealed a reduction in the length of the posterior lobe, while it was wider (Fig. 2F), suggesting that newly delaminated neuroblasts could not extend posteriorly when the pioneer axon was missing. In some extreme cases, the posterior lobe was absent (Fig. 2F). Altogether, the data indicates that pioneers are fundamental for neuroblast migration and ultimately organizing the lobular shape of the SAG.

We also analyzed the extension of central projections, since it occurs simultaneously. Small central axons began to grow at 20 hpf, initially defasciculated and emitting dynamic branches (Fig. 2G, red arrowheads). Progressively, the central pLL axons fasciculated into a bundle, while the pLL ganglion displaced posteriorly (Fig. 2G). The SAG and LL central projections entered the hindbrain, turned and grew in parallel to the hindbrain border and lateral to the trigeminal bundle. The entry points of central projections to the hindbrain are referred to as the anterior and posterior entry points (aEP and pEP, respectively, Fig. 2G). Three separate bundles could be observed in an orthogonal slice of the sensory axons extended along the hindbrain; the trigeminal (Tg), the aLL- SAG and the pLL bundle, from medial to lateral as described previously (Fig. 2G; Zecca et al., 2015).

Overall, our live imaging of SAG and LL extending axons to their targets, reveal new information on how the mechanosensory circuit develops: i) SAG pioneer axons are the first to contact HC, but also establish interactions with pLL central projections; ii) peripheral and central projections progressively fasciculate into a bundle; and iii) otic neuroblasts use the SAG pioneer axons to migrate and establish the posterior lobe of the SAG (Fig. 2H).

### Cxcl14 is an early chemokine dynamically expressed in first-born HCs and hindbrain entry points

When screening factors that might direct pioneer SAG axons, we found that the chemokine *cxcl14* was one of the earliest molecules expressed at the first-born HC of the inner ear but not HCs from LL. The pattern of *cxcl14* expression in the inner ear is dynamic and complex. At 24 hpf, *cxcl14* was restricted to the prospective anterior and posterior macula domains (am and pm, respectively) of the OV (Fig. 3A). The expression coincided with the first HCs labeled by Tg(*brn3c:eGFP) ^s356t^* (Fig. 3A’, magenta and black arrowheads) that are already innervated by axons at 26 hpf (Fig.3B, B’). By 32 hpf, *cxcl14* was expressed in three and two HCs of the am and pm, respectively (Fig. 3C-C’’’, black arrowheads). Interestingly, *cxcl14* also appeared in two spots adjacent to the OV and to the hindbrain (Fig. 3D, D’’, magenta and white arrowheads). Analysis of *cxcl14* expression combined with axonal staining at 36 hpf, revealed tha*t cxcl14* expression in these two spots corresponded to the hindbrain entry points of SAG and LL central projections to the hindbrain (Fig. 3E, magenta and white arrowheads). At 36 hpf, *cxcl14* was still expressed in the am and the two entry points but, strikingly, expression in the pm only remained in one or two cells (Fig. 3E’ and E’’, black arrowheads). Later, at 48 hpf and 72 hpf, *cxcl14* expression was only present in the cristae, as these sensory domains generate HCs later (Fig. 3F-G), while the expression in maculae HCs disappeared.

**Figure 3.**
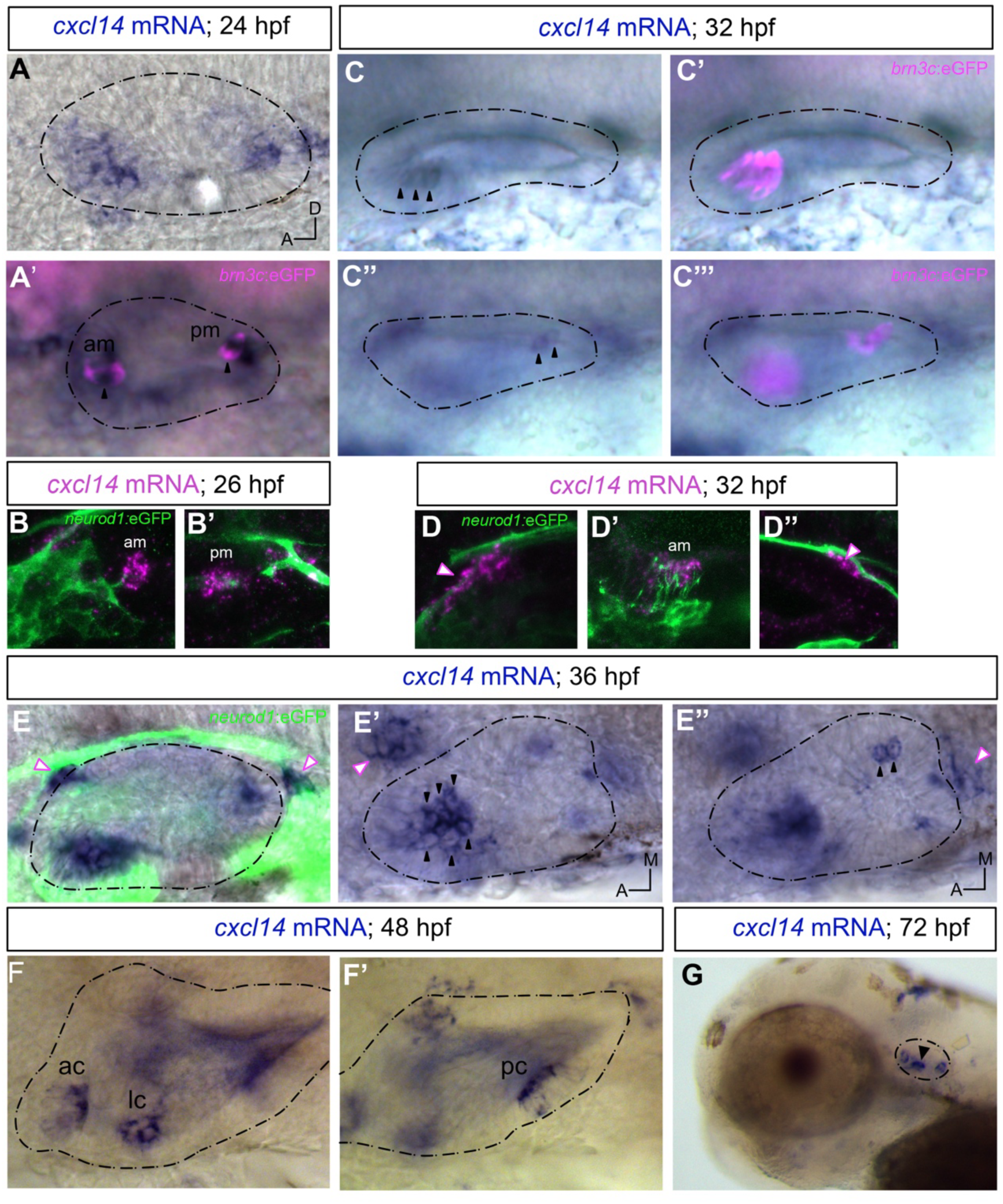
*cxcl14* is expressed in inner ear HCs and hindbrain entry points. Chromogenic in situ hybridization (ISH) of *cxcl14* mRNA at different timepoints. (A) *cxcl14* is expressed in tether HCs of the am, pm at 24 hpf (black arrowheads). HCs are visualized in magenta (A’). (B) Fluorescent ISH at 26hpf with neurons labelled in green showing the am and the pm and how *cxcl14*^+^ HC are innervated by pioneer axons. (C) Images at 32 hpf with the focus on the anterior region of the OV (C, C’) and the posterior region (C’’, C’’’, HC in magenta). HCs expressing *cxcl14* are marked with black arrowheads. (D) Fluorescent ISH at 32hpf with neurons labelled in green showing expression at the anterior entry point (D, magenta and white arrowheads), am (D’) and posterior entry point (D’’, magenta and white arrowheads). (E) Images at 36 hpf with neurons and axons labelled in green (E). In (E’), the anterior region of the OV is focused and in (E’’), the posterior region, showing am and pm, respectively. Hindbrain entry points are marked by magenta and white arrowheads. HCs with *cxcl14* expression are marked by black arrowheads. (F) *cxcl14* expression at 48 hpf in the ac, lc (F) and pc (F’). (E) *cxcl14* expression at 72 hpf. Black dotted lines depict the OV silhouette. ac: anterior crista; lc: lateral crista; pc: posterior crista; am: anterior macula; pm: posterior macula. Neurons are labelled through TgBAC(*neurod1:eGFP*)^nl1^ and HC through Tg(*brn3c:eGFP*) ^s356t^.

Hence, the expression pattern of *cxcl14* chemokine is complex and highly dynamic, linked at sites and timepoints of pioneer axonal targeting; while disappearing once axons reach the target sites.

### Depletion of *cxcl14* results in axon growth and targeting defects in SAG pioneer axons and hindbrain projecting axons

The expression pattern of *cxcl14* in nascent HCs when afferent-HC connections are established, together with expression at the regions where central axons grow into the hindbrain, suggests a role of Cxcl14 in axon guidance and growth. To assess this, we performed functional analysis in *cxcl14* F_0_ CRISPR crispants (see Material and Methods). Compared to control embryos, SAG pioneer axons displayed navigation and targeting errors. A significant number of SAG axons did not follow the usual growth path (58.3%) and incorrectly targeted the posterior prosensory domain (41.7%), while we observed no errors in control cases (Fig. 4A, F and H). In control embryos, the growth of pioneer axons followed a horizontal and straight path towards the posterior prosensory domain (Fig.4A, C). However, in crispant *cxcl14* embryos, growing axons explored a larger territory and grew following a tortuous path, even touching the hindbrain or growing along the central trigeminal axons (Fig. 4B, C; Video 4,5). Thus, the angle and direction of growth in *cxcl14* crispants were wider than in controls (Fig 4D, E). Misrouted axons never grew towards the underlying mesenchyme. The growth of SAG pioneer axons in crispant embryos was also slower and showed more dynamic behavior, as axons tended to elongate and retract constantly, exhibiting unstable pathfinding behavior (Fig. 4G). The exploration of more territory can also be explained due to a fasciculation impediment in *cxcl14* depleted embryos (only 30% of crispant embryos showed axon fasciculation, while 90% of controls exhibited fasciculated axons) (Fig.4E, H).

**Figure 4.**
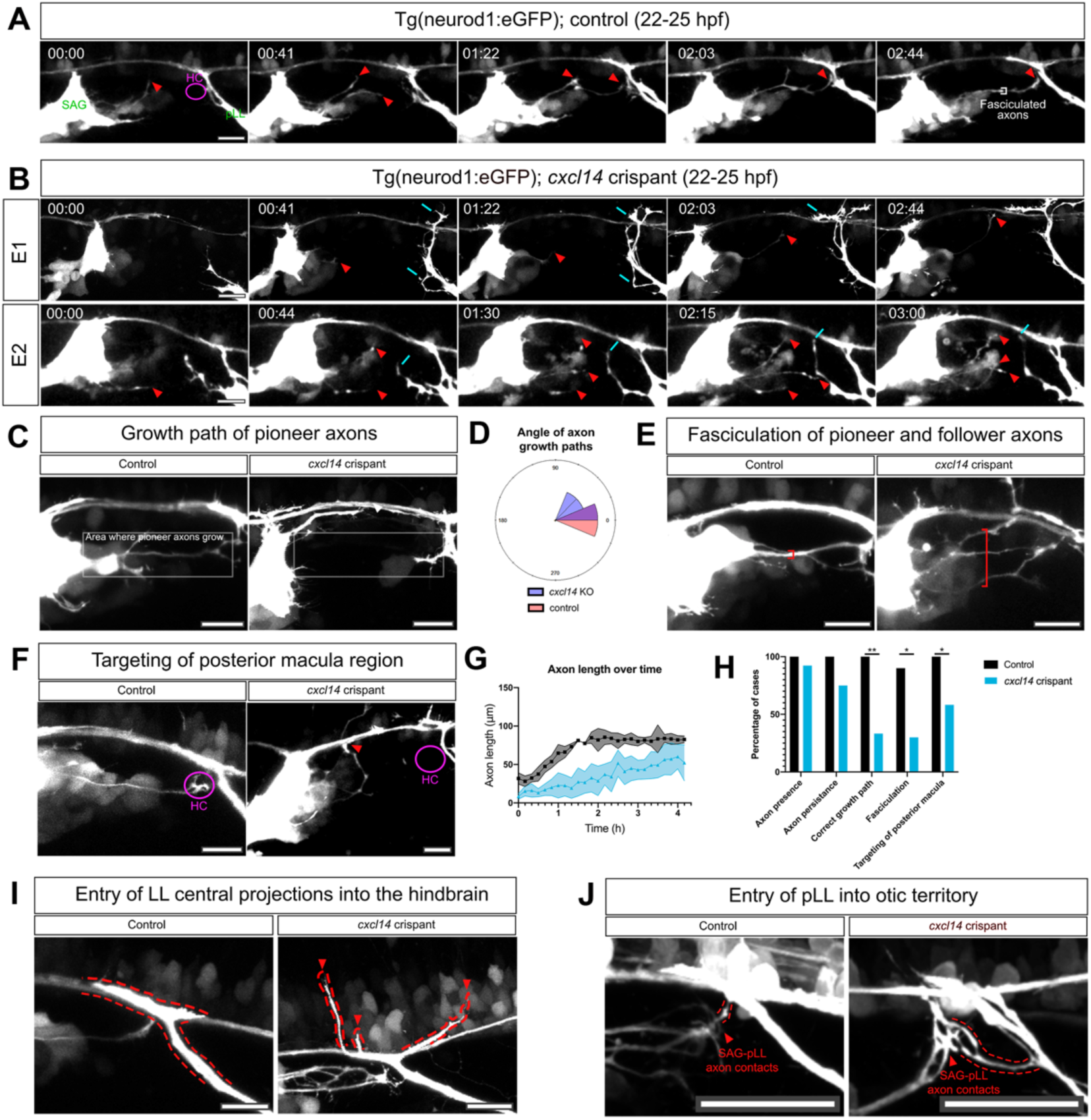
*cxcl14* depletion impairs axon growth, targeting and fasciculation. (A,B) Stills of a time series showing the growth dynamics of SAG pioneer axons in a control (A) and two representative examples of *cxcl14* crispants (B) from 22 to 25 hpf. Arrowheads mark the growth cone of axons. Magenta circle marks the theoretical position of the first HCs. White square bracket indicates fasciculated axons. pLL axons appear defasciculated and send projections to otic territory (blue arrows). (C) Confocal max projections of a control and a *cxcl14* crispant. Dotted-line rectangle represents the area in which SAG pioneer axons grow in control embryos. In *cxcl14* crispant embryo pioneers axons grow outside of this area. (D) Rose plot representing the angle of SAG pioneer axon growth path (control vs *cxcl14* crispant). N = 5 each. (E, F) Representative confocal max projections of a control and a *cxcl14* crispant SAG pioneer axon fasciculation (E) and targeting (F). Red square brackets in e indicate the vertical dispersion of the group of axons, from most dorsal to most medial. Magenta circles in F represent the theoretical position of newborn HC. Red arrowhead indicates the growth cone of the crispant example. (G) Line plot representing the SAG pioneer axon length over time of controls (grey) vs *cxcl14* crispant (light blue). N = 5 each. Dispersion represents SD. (H) Bar plot representing the percentage of cases (controls vs *cxcl14* crispant embryos) that comply with the categories: axon presence, axon persistence, correct growth path, fasciculation and targeting of posterior macula. N = 10 controls / 12 cxcl14 crispants. (I,J) Representative confocal max projections of a control and a *cxcl14* crispant pLL axon entrance to the hindbrain (I) and to otic territory (J) (highlighted with red dotted lines). Arrowheads in I mark axons that grow dorsally in *cxcl14* crispant. Arrowheads in J mark the contact between SAG pioneer axon and pLL axon. Scale bars = 20 µm. Tg(*neurod1:eGFP*)^nl1^ is used to label sensory neurons and axons.

We then analyzed the phenotypes from pLL axons projecting to the hindbrain. In control embryos, all central axons entered the hindbrain, fasciculated, and turned to position themselves in parallel and lateral to the trigeminal axons. However, in crispant embryos, pLL axons failed to enter the hindbrain (15,4%) and only 9% of the pLL central projections entered correctly. They exhibited growth and turning errors, growing perpendicular to the anteroposterior axis, in a dorsal direction (Fig. 4I). Also, the contacts that were established between the SAG pioneer axons and the pLL axons happened in both control and crispants, but contacts were longer in crispants, with more axons from the pLL invading the otic (Fig. 4J).

It has been reported that *cxcl14* expression depends on *atoh1b* (Ezhkova *et al*., 2023). Thus, we knocked down *atoh1b* by injecting a morpholino and examined SAG pioneer axons. In morphant *atoh1b* embryos, most SAG pioneer axons extended posteriorly, and axons lost their characteristic axon swelling in the HC contacting region compared to wild-type embryos and grew beyond the otic vesicle (Fig. Suppl. 2, arrowhead in inset). As in *atoh1b* morphants, *cxcl14* is depleted from the inner ear but not from the entry points, SAG axons could still extend posteriorly, presumably due to the non-otic posterior source of Cxcl14. As expected, in *atoh1b* morphants, pLL central projections did not present any fasciculation nor turning phenotype (Fig. Suppl. 2). The experiment indicates that pioneer axon initial growth is independent of HC specification, but *atoh1b* and *cxcl14* are required for proper targeting of the growth cone to HCs and preventing long-lasting connections with other growing axons. Hence, early-born sensory cells act as spatial organizers for circuit assembly, rather than passive targets of innervation.

### Ectopic expression of Cxcl14 reroutes SAG afferent axons and lateral line central projections

To test whether Cxcl14 can attract and reroute axons, we ectopically expressed Cxcl14 by injecting the *hsp70I*:*cxcl14-IRES-CAAX-GFP* plasmid, heat-shocking embryos at 16 hpf and mounted for live imaging (Fig. 5A). We selected embryos in which ectopic *cxcl14*-expressing cells were located near the OV (Fig. 5C). We observed, in 72% of the cases (N=11), axons directed towards cells expressing Cxcl14 and, in some cases, contacting directly with them (Fig. 5C, white arrowheads). We could see that the SAG pioneer axons and the central projections of the pLL were the ones rerouting, compared to axons in controls (Fig. 5B and C). Some axons abandoned the fascicle directed to the hindbrain and grew towards cells expressing exogenous Cxcl14 (N=4, Fig. 5C). In one case, we could even observe a SAG pioneer axon entering the hindbrain to target cells expressing Cxcl14 (Fig. 5C). Rerouting of axons was only observed when the ectopic source of Cxcl14 was relatively nearby, possibly due to a short-range activity or other tissue constraints. Based on these data, we concluded that Cxcl14 acts as an attractive signal for SAG pioneer and pLL central axons.

**Fig. 5.**
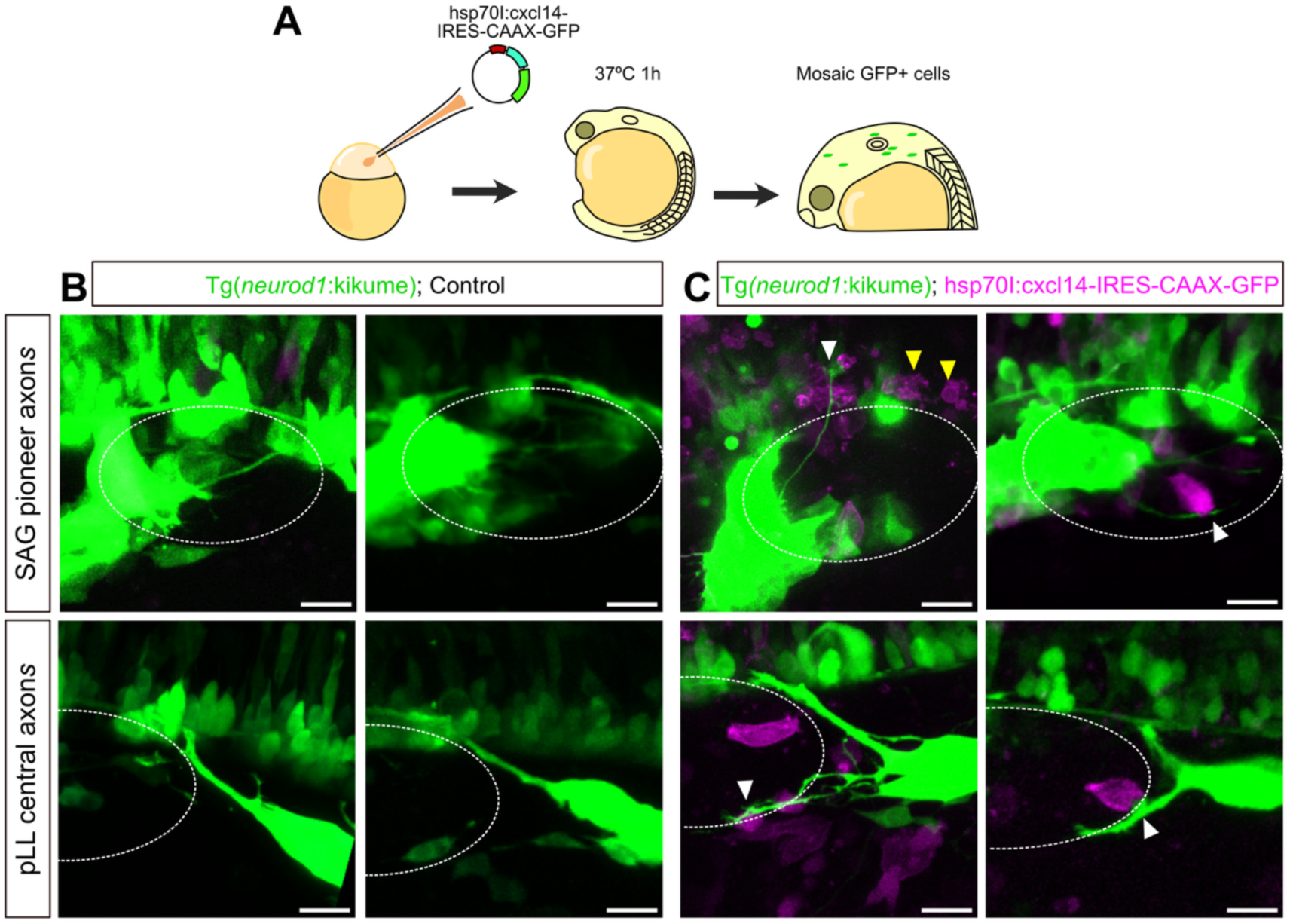
Ectopic expression of *cxcl14* reroutes axons. (A) Schematic representation depicting the generation of a transgenic larvae that expresses *cxcl14* ectopically under the control of Hsp70I promoter. (B-C’’’) Confocal max projections of control embryos focusing on SAG axons (B-B’) and pLL axons (C-C’). (B’-C’’’) Confocal max projections of embryos injected with hsp70I:cxcl14;IRES:CAAX-GFP plasmid and heat-shocked to ectopically express *cxcl14* (in magenta). White arrowheads indicate rerouted axons towards plasmid-expressing cells. Yellow arrowheads indicate dying cells. Scale bars = 10 µm. Neurons are visualized by photoconversion of Tg(*neurod1:kikume*) to differentiate between kikume and GFP from the plasmid.

### Hair cell activity in cristae is compromised and Otoferlin is mislocalized in *cxcl14* **crispant embryos**

The action of chemokines has been shown to be paracrine, as well as autocrine, and in the central nervous system, Cxcl14 regulates calcium and GABA release (Banisadr *et al*., 2011; Iannone *et al*., 2024). Following this rationale, we assessed whether loss of *cxcl14* has any effect on HC function. HC activity was assessed with a 4-Di-2-ASP vital mitochondrial dye uptake assay, which is based on the entrance of this fluorophore through active mechanotransduction channels (Faucherre *et al*., 2009; Rubbini *et al*., 2015). While the total number of HC (quantified by GFP labelling) did not show any significant change in *cxcl14* crispants, the ratio of active HCs/ total HCs (4-Di-2-ASP+/GFP+) was significantly reduced in ac and lc (Fig. 6A-C), and to a lesser extend in pc (Fig.6D). As internal control, we also assessed the activity in HCs of the neuromasts where *cxcl14* is not expressed and no significant differences were found in neuromast HCs (Fig. 6A, E). The lack of HC activity in *cxcl14*-deficient embryos might indicate a defect in mechanotransducing channels or a lack of maturation of HCs.

**Fig. 6.**
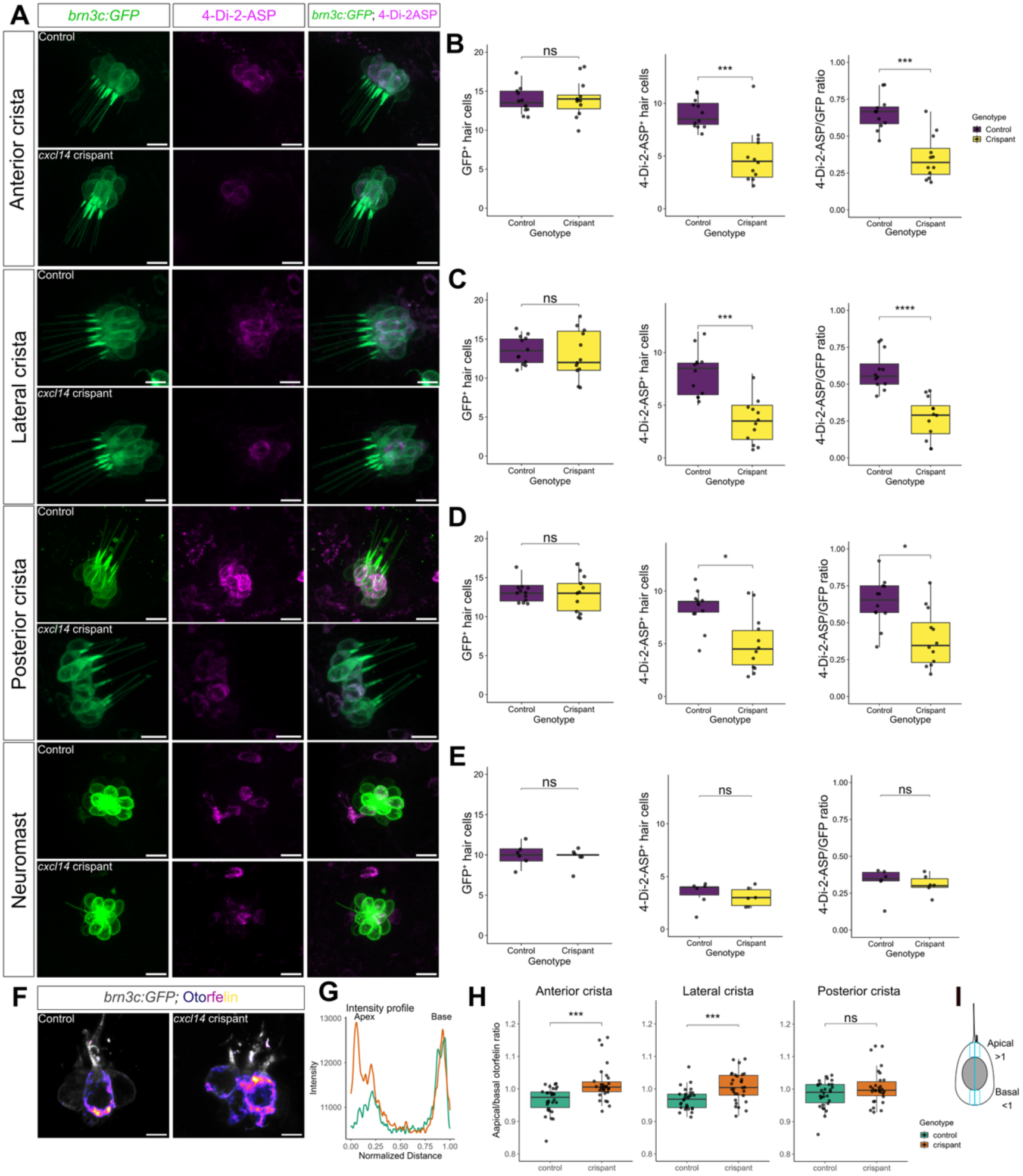
c*xcl14* depletion has an effect in HC activity and Otoferlin distribution. (A) Representative confocal images of AC, LC and PC hair cells labelled by Tg(*brn3c:GFP*) *^s356t^* (green) and 4-Di-2-ASP labelling (magenta) at 72 hpf in control and *cxcl14* crispants. Scale bar = 5 µm. (B, C, D, E) Boxplots showing the quantification of GFP+ HC, 4-Di-2-ASP+ HC and the ratio 4-Di-2-ASP/GFP+. **GFP+ HCs**: ac= 13.67 ± 1.72 (controls) and 13.08 ± 3.06 (crispants), p=0.53; lc= 13.83 ± 1.47 and 14.08 ± 2.35, p=0.91; pc = 13.17 ± 1.19 and 12.92 ± 2.39, p=0.52; neuromast = 10 ± 1.29 and 9.67 ± 1.24, p=0.86. **4-Di-2-ASP+ HCs**: ac= 8.08 ± 1.72 (controls) and 3.5 ± 3.06 (crispants), ***p=0.00067; lc = 9 ± 1.37 and 5.08 ± 2.35, ***p=0.00018; pc = 8.33 ± 1.19 and 5.08 ± 2.39, *p=0.01; neuromas = 3.33 ± 1.1 and 3 ± 0.8, p=0.44. **Ratio:** ac = 0.59 ± 0.13 (controls) and 0.27 ± 0.14 (crispants), ***p=0.00033; lc = 0.66 ± 0.11 and 0.35 ± 0.15, ****p=0.000067; pc = 0.64 ± 0.16 and 0.39 ± 0.19, *p=0.01; neuromast = 0.33 ± 0.1 and 0.31 ± 0.07, p=0.42. N = 12 controls / 12 *cxcl14* crispants from 2 independent experiments. Each dot represents one embryo. (F) Representative z-slice confocal images of HC labelled in grey and immunostained for Otoferlin (fire LUT). Scale bar = 5 µm. (G) Example of intensity profile of Otoferlin signal obtained from drawing a line from the apex to the base of HC in control (turquoise) and *cxcl14* crispant (orange). (H) Boxplots of the ratio between apical/basal Otoferlin signal in control (turquoise) and *cxcl14* crispants (orange). Each dot represents one cell (n = 30 cells from N = 5 embryos of each group). (I) Schematic representation of how the measurement of Otoferlin signal was performed in a HC. A ratio <1 = more apical expression; ratio >1 = more basal expression. Mann–Whitney test was performed in all comparisons.

Following our analysis of HC activity, we evaluated the localization of Otoferlin, a calcium sensing protein that mediates synaptic vesicle release specifically in the ribbon synapses of HCs. By plotting the intensity profile of expression in a vertical line from the apical to the basal part of HCs and calculating the ratio of expression in apical/basal (Fig. 6F-I), we found that, in the ac and lc of control embryos, Otoferlin was preferentially localized in the basal area of HC where synaptic vesicles concentrate (ratio <1, ac mean ratio = 0.96, lc mean ratio = 0.97), while in *cxcl14* crispants, Otoferlin is more evenly distributed in apical and basal regions (ratio = 1.00, ac mean ratio = 1.02, lc = 1.01) (Fig. 6H).

Thus, these results demonstrate that Cxcl14 is necessary for HC activation and synaptic organization, potentially causing subsequent failure in the transmission of mechanosensory information.

### Cxcl14 regulates the complexity of afferent axonal arbors in cristae

Then, we analyzed afferent SAG axons arborization in the cristae where axonal terminals are well separated from somas, facilitating their imaging and HCs present reduced activity. We traced and 3D-reconstructed axonal terminal arborizations in anterior, lateral and posterior cristae (ac, lc and pc, respectively) (Fig. 7A-F’’) using the Single Neurite Trace (SNT) plugin in Fiji. In controls, fasciculated bundles were sent from somas of the SAG to each specific crista, and when they reached them, axons turned at an angle higher than 90° and ramified to innervate each hair cell.

**Fig. 7.**
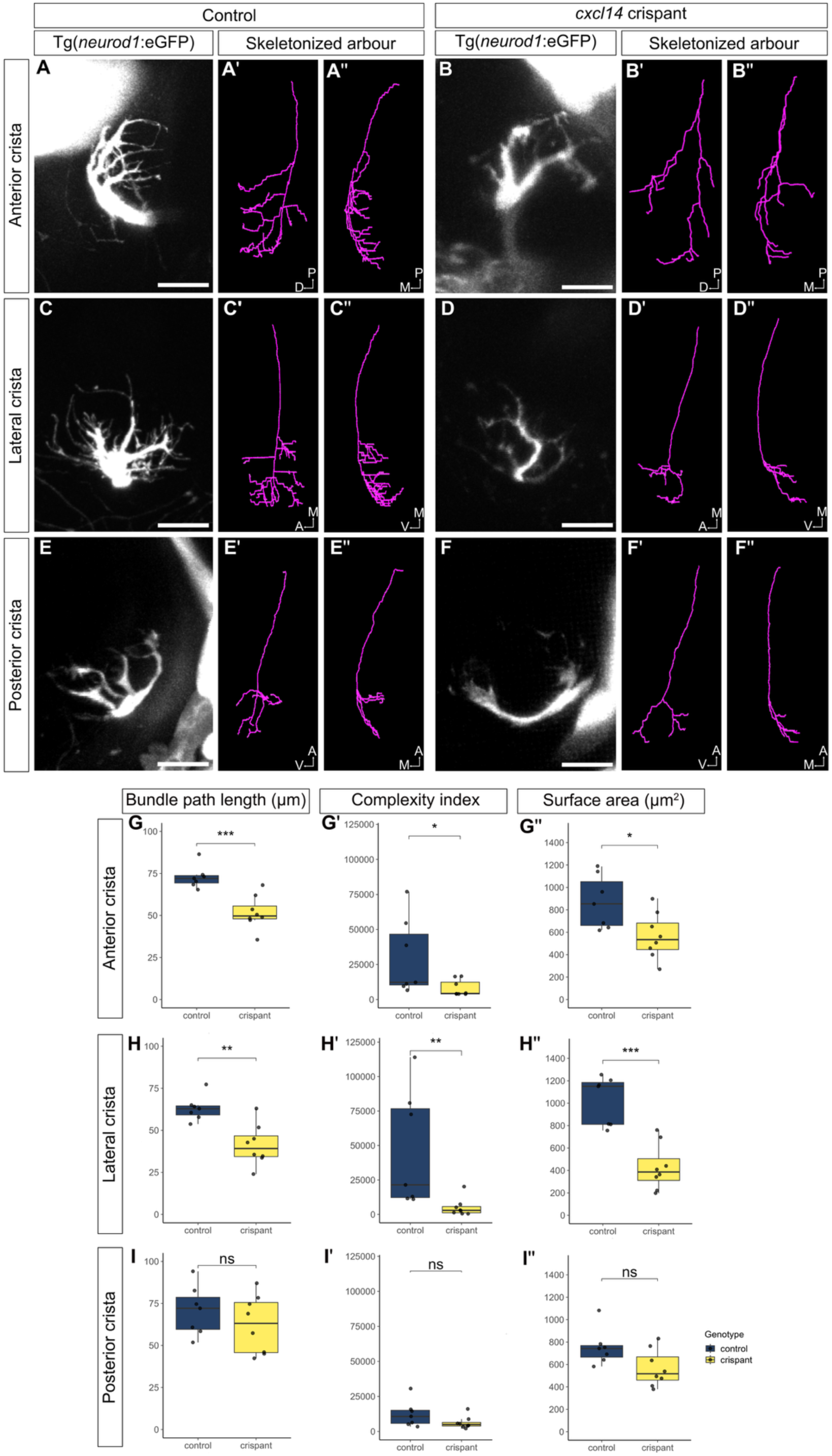
*Cxcl14* depletion causes a reduction in axonal arbour morphology complexity in innervation sites of cristae. (A-F’) Representative examples of control and *cxcl14* crispant axonal arborizations in anterior crista, lateral crista and posterior crista innervation sites. Each panel depicts the confocal max projection of axonal terminations visualized by TgBAC(*neurod1:eGFP*)^nl1^ and a skeletonized reconstruction of the axon from two different views (magenta). Scale bars = 10 µm. (G-I’’) Boxplots showing the quantification of axon bundle path length (µm) (anterior crista = 72.8 ± 2.55 for controls and 51.0 ± 3.48 for *cxcl14* crispants, ***p=0.0006; lateral crista = 63 ± 2.79 and 41.3 ± 4.29, **p=0.004; posterior crista = 70.6 ± 5.59 and 62.4 ± 6.05, p=0.4), and axonal arbour complexity index (ac = 29979 ± 10367 for controls and 8084 ± 2031 for *cxcl14* crispants, lc = 46329 ± 15924 and 5112 ± 2305, **p=0.002; pc = 12354 ± 3515 and 6230 ± 1571, p=0.15) and surface area (µm^2^) (ac = 870 ± 89 for controls and 567 ± 72.3 for *cxcl14* crispants,*p=0.029; lc = 1022 ± 82 and 429 ± 72, ***p=0.0006; pc = 754 ± 60.7 and 567 ± 57.8, p=0.07) in control (dark blue) and *cxcl14* crispant (yellow) embryos for each crista. N = 7 controls / 8 *cxcl14* crispants from 2 independent experiments. Mann–Whitney test was performed in all comparisons.

In crispant embryos, axon bundles were still growing towards the cristae. Nevertheless, the overall ramifications were composed of fewer branches. We then quantified several parameters of axonal terminals to assess their complexity (Fig. 7G-I’’). Regarding the ac, in 50% of crispant embryos (N=8), branches were misrouted, resulting in some axons defasciculated from the bundle and the fasciculated bundle was shorter (Fig. 7B, B’’). In the case of lc, there were cases in which axons were hard to distinguish, as they were very few and thinner than in controls (data not shown). The length of the fasciculated bundle was also reduced in crispant embryos (Fig.7G-I), compared to controls (N=7), and the axonal complexity index, i.e. how intricate an axonal arbor is, and axonal surface area were dramatically reduced, indicative of a reduction in innervation capability (Fig. 7G-I’’). Surprisingly, pc was less affectated than the other cristae, showing a tendency of a lower complexity index and smaller surface area; although, these differences were not statistically significant (Fig.7I-I’’). Together, these results show that Cxcl14 is involved in axonal arborization in cristae and that the lack of the chemokine results in less complex branches covering a smaller area.

### The vestibuloacoustic response is impaired in *cxcl14* crispant embryos

To test whether the phenotypes observed at a cell-resolution level were affecting the overall vestibuloacoustic system’s responses, we performed a behavioral test using the DanioVision observer chamber. We tested the startle response in front of a tapping stimulus on 5 days post fertilization (dpf) *cxcl14* crispant and control larvae. The startle response is an escape mechanism that appears from 5dpf onwards, characterized by a rapid C-bend of the body followed by swimming. It is mainly mediated by the vestibuloacoustic system, which senses vibrations; Mauthner cells integrate the sensory input and initiate the motor response.

Generally, *cxcl14* crispant larvae moved less than their counterpart control siblings (Fig. 8A, A’, A’’). We visually analyzed the tracking recording and observed that after each tapping, a smaller number of *cxcl14* crispant larvae showed a startle response (the average value of *cxcl14* crispant larvae that moved after tapping was 4.2 out of 22 larvae ± 2.39) in comparison to control larvae (14.9 out of 23 larvae ± 2.56) (Fig 8B). The intensity of the startle response was also diminished in *cxcl14* crispants (Fig. 8B’, B’’), which was calculated as the average of individual startle response corrected by the baseline activity of each larva at each tapping stimulus (Fig 8B’’). This resulted in a lower distance moved by *cxcl14* crispants than by controls. Although not shown in the tracking, we could observe that *cxcl14* crispant larvae fell laterally when they were not swimming, a sign of lacking stationary balance. Therefore, these results show that the lack of *cxcl14* has an effect in vestibuloacoustic function in 5dpf larvae.

**Fig. 8.**
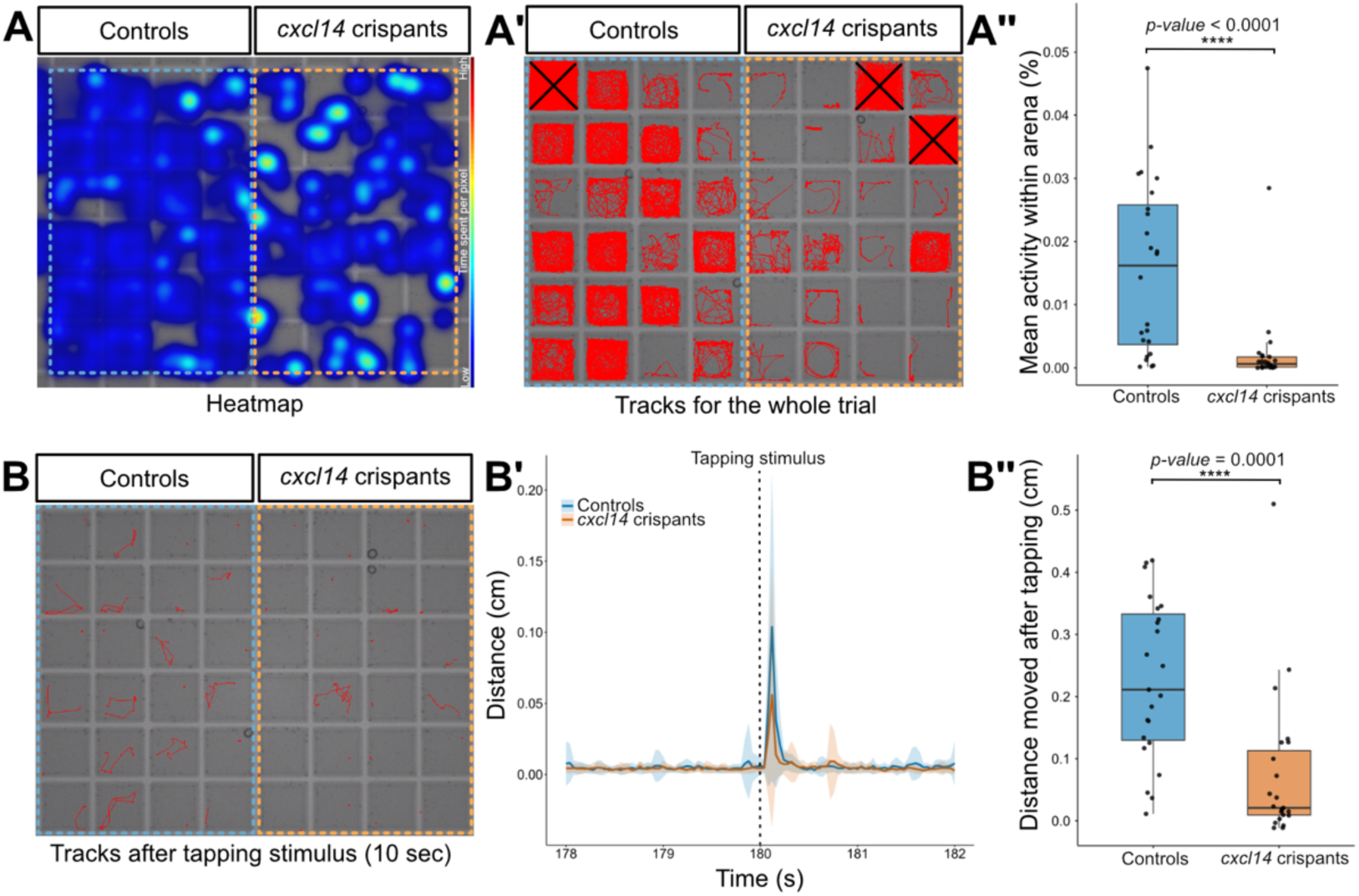
Startle response after acoustic stimuli is reduced in *cxcl14* crispant 5dpf larvae. (A) Heatmap of the total time spent per pixel during the whole recording. Colder colours in the scale represent a lower amount of time that the larvae have stayed on a pixel, while hotter colours represent a higher amount of time. (A’) Tracking of the path (red) followed by larvae on their corresponding wells during the entire experiment. Wells marked with an X were discarded for further analysis, due to faulty tracking. (A’’) Boxplot representing the mean individual activity of control (blue) and *cxcl14* crispant (orange) larvae, represented as the % of the total area of the arena, which corresponds to each well. Individual values are represented as dots. Control group = 0.016 ± 0.013, *cxcl14* crispants = 0.002 ± 0.006. p-value = 0.0000029. (B) Tracking of the path (red) followed by larvae in the period of 10 sec after a tapping stimulus. (B’) Example of the distance moved by larvae pre and post-stimulus. Line plot of the distance moved by the larvae two seconds pre-stimulus and two seconds post-tapping. The stimulus is marked with a dotted line at the second 180. (B’’) Startle response strength. Boxplot representing the individual average of the distance moved one second after tapping stimuli normalized by the baseline of each larva (dots represent individual values for each larva). Control group = 0.227 ± 0.026; *cxcl14* crispants = 0.074 ± 0.036. p-value = 0.00012. Values are given as mean ± S.D. Mann-Whitney test was performed for the activity within arena comparisons, and a linear mixed-effects model was used for the startle response strength comparisons.

## Discussion

Our findings uncover two general principles of broad relevance. First, peripheral sensory circuits utilize pioneer-based cellular scaffolds analogous to those described in the CNS, challenging the prevailing view that inner ear circuitry derives exclusively from otic-derived neurons. Second, our work reveals chemokines as bona fide regulators of axon navigation during development, extending their functional repertoire beyond immune and inflammatory contexts. In particular, we reveal a new role of the little studied Cxcl14 chemokine in directed axonal growth and HC activity.

In mouse inner ear, several guiding molecules, have been proposed to guide auditory and vestibular afferents to the hair cells located in the sensory epithelium at late developmental stages (Hashimoto and Heinrich, 1997; Fritzsch *et al*., 2005; Kim *et al*., 2016; Jung *et al*., 2019; Cantu-Guerra *et al*., 2023). However, early developing axons extend to sensory regions before hair cells are fully differentiated (Hemond and Morest, 1992), suggesting that early secreted otic vesicle trophic factors fulfill this guidance role. In particular, the chemokine MIF was found to promote the growth of chick and mouse SAG axons *in vitro* (Bianchi *et al*., 2005), but its expression was not found in the developing sensory patches in vivo. Thus, how the first axons are directed to that specific region to innervate the HCs have not been resolved.

Here we show that pioneer axons are extended by a population of pioneer cells of non-otic origin sharing markers with aLL progenitors. Both aLL and otic progenitors derive from a common OEPD domain expressing *pax2* and *pax8*, and although aLL and otic progenitors segregate over time, it is plausible that at early stages an uncommitted population of cells serves as pioneers for SAG and aLL ganglion development. Whilst the role of pioneers in pLL has been investigated (Gompel, Dambly-Chaudière and Ghysen, 2001; Pujol-Martí and López-Schier, 2013; Woodruff *et al*., 2025), this is the first report of a role of pioneer axons in inner ear development, showing that pioneer axons serve as scaffolds for neuroblast migration and organization of SAG lobes.

The use of the zebrafish has also permitted us to image at high spatiotemporal resolution how pioneer SAG axons extend towards HCs. As shown in other contexts (Ma and Tessier-Lavigne, 2007; Lewis, Courchet and Polleux, 2013; Kalil and Dent, 2014), axons emitted dynamic branches that eventually were retracted, but SAG pioneer axons and LL axons established transient contacts that were maintained for longer time periods. It has recently been proposed that axon-axon interactions help to establish a correct somatosensory map in *Drosophila* (Galindo *et al*., 2023). When cIV neurons were ablated, axons of cIII invaded new territories, suggesting that transient axon-axon interactions might have an inhibitory effect on their growth and targeting area. Therefore, it is plausible that transient SAG and LL axon-axon interactions are a developmental mechanism to finally segregate each circuitry, that develops in proximity. Thus, the correct growth and targeting of SAG pioneer axons seems fundamental for the precise contact with pLL axons, allowing the posterior segregation of the two distinct axons.

The highly dynamic and specific expression pattern of *cxcl14* in first-born HCs provides cues for its function. Interestingly, its expression in the pm was restricted to only the first two HCs that differentiated in this sensory patch, suggesting a difference between the two firstly differentiated HCs and the rest. We hypothesize that the innervation of the first-born HCs is different from the later-born HCs, thus *cxcl14* serves to promote the posterior navigation of SAG pioneer axons to correctly target the first HCs of the pm. On the contrary, the later-born HCs of the pm are innervated by neuroblasts from the posterior lobe of the SAG, whose somas are in close contact with the macula and their axons do not navigate nor extend long distances to find the HCs, as they are already oriented towards the base of HCs (Fig. 9). Regarding the am, HCs are located far from the SAG somas, thus axons grow beneath the inner ear and turn dorsally, to innervate the HCs of the am. Hence, Cxcl14 at the am could be guiding the first axons and might not be necessary for the follower axons, thus explaining its downregulation. Supporting our hypothesis, axons targeting the cristae also navigated long distances. Expression in cristae was kept at least high until at least 5 dpf (Daniocell database and data not shown), which can be explained to the morphogenetic displacement of cristae that would require axons to grow more than the maculae counterparts. The observed loss of directional growth of SAG pioneer axons to posterior HCs in *cxcl14-*depleted embryos, together with the reduction of axonal arborization complexity and area covered by axons, aligns with our hypothesis. Furthermore, in overexpression experiments, some axons were misrouted and extended to ectopic *cxcl14*-expressing cells, suggesting a role of the chemokine in guiding certain axons.

**Fig. 9.**
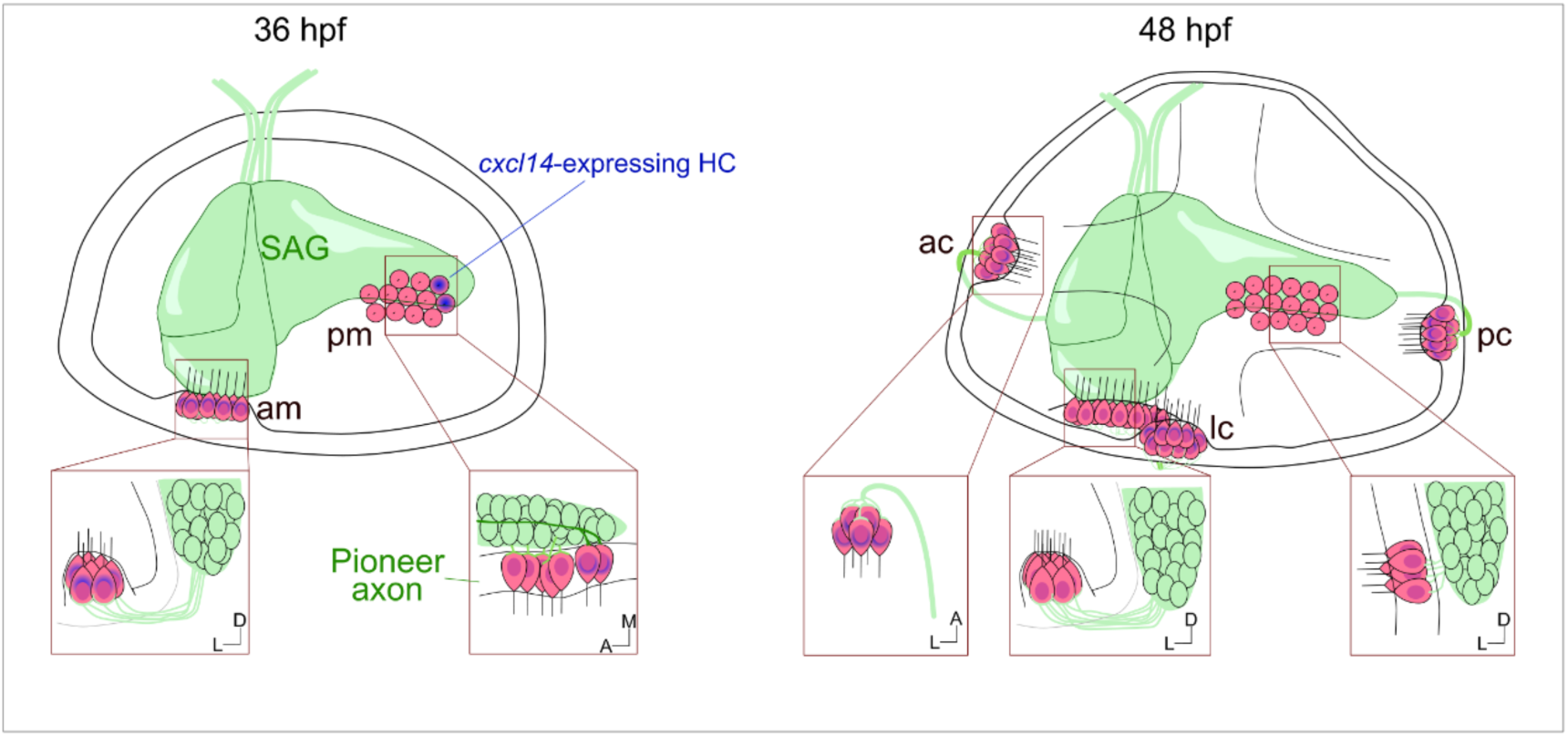
Schematic representation of the spatial relationship between neurons and HCs. (Schematic drawings of a 36 and 48 hpf OV, depicting SAG neurons and axons (green) and HCs (pink). *Cxcl14*-expressing HCs are depicted in blue.

As the expression of *cxcl14* in entry points initiated concomitantly with the extension and fasciculation of central axons, and in *cxcl14*-depleted embryos, central projections failed to correctly enter the hindbrain, displayed navigation, fasciculation and turning errors, we also propose that *cxcl14* regulates central projections growth to the hindbrain. It would be interesting to assess whether in other vertebrates *cxcl14* is also present in entry sites of central projections and regulates these projections. Finally, the nature of the cells expressing *cxcl14* in the entry points is still elusive. They might have a neural crest origin, as neural crest cells migrate in two streams adjacent to the otic placode and, in the dorsal root ganglion, neural crest-derived boundary cap cells regulate the entry and exit of sensory and motor axons (An, Luo and Henion, 2002). Further studies should investigate this question.

Despite the effect of the chemokine in axonal growth and targeting, additional roles in HC physiology cannot be excluded. We found reduced HC activity and Otoferlin was not properly distributed towards the base of HCs in *cxcl14*-depleted embryos, suggesting a disruption of synapse vesicle trafficking. The HC phenotype could be a direct effect of the lack of *cxcl14* or an indirect effect from an improper axonal innervation. In mouse, afferent contacts preceded presynaptic ribbon attachment in the base of HCs (Michanski *et al*., 2019), suggesting a possible correlation between innervation and HC synaptic activation. Nevertheless, studies in zebrafish lateral line neuromasts demonstrated that HCs were active in larvae that lacked innervation (López-Schier *et al*., 2004). Therefore, c*xcl14* could be affecting both HC activity and axon growth and targeting. In this regard, recent work in the somatosensory cortex, revealed that Cxcl14 is required for regulating SBC axonal morphology and excitability (Iannone *et al*., 2024).

As both HCs and axonal innervation were affected by the lack of *cxcl14*, we tested if the implication of the chemokine was translated as a general malfunctioning of the vestibuloacoustic system. Indeed, we found an impairment in the stationary balance of 5dpf *cxcl14* crispant larvae, together with a reduced startle response when larvae were submitted to acoustic stimuli. The reduction in general locomotor activity could be explained due to the lack of stationary balance, locomotor affectations due to vestibular defects (Bruce B. Riley, 2000; Kim *et al*., 2014) or other locomotor impairments non-related. However, taking into consideration that the startle response was reduced, the results suggest that larvae lacking *cxcl14* could not respond to vibrational stimuli as much as control larvae, meaning that the correct function of the statoacoustic sensory system relies on the chemokine *cxcl14*.

Due to the evolutionary conservation of *cxcl14* in inner ear HCs in all vertebrates analysed so far, our work in zebrafish sets the basis for future functional studies in other vertebrates. Interestingly, recent scRNA-seq datasets identified *cxcl14* to be enriched in mouse and human utricles (Wang *et al*., 2024), in regenerating type II HC in mouse and human utricles (Luca *et al*., 2025), in tall HC of the postnatal day 7 chick cochlea and in chick regenerating utricle HC (Ku *et al*., 2014; Janesick *et al*., 2022). As our work provides functional evidence for the requirement of Cxcl14 in HC activity and setting the pioneer neural circuit, we speculate that Cxcl14 induction in regenerating HCs might be necessary for reinnervation of HCs after injury. Nevertheless, further studies are needed to confirm this hypothesis in regenerating tissues.

Ultimately, these findings fundamentally reshape our understanding of peripheral neurodevelopment by demonstrating that inner ear circuit assembly relies on pioneer axonal scaffolds and expands the functional repertoire of chemokines into essential developmental axon guidance cues.

## Materials and methods

### Fish maintenance and husbandry of transgenic lines

Zebrafish embryos and adults were maintained and handled according to standard procedures at the aquatic facility of the Parc de Recerca Biomèdica de Barcelona (PRBB), in compliance with the guidelines of the European Community Directive and the Spanish legislation for the experimental use of and as previously described (Westerfield M., 2000). Stable transgenic lines were kept by means of alternate outcross with WT (AB/Tü) and incross, generation after generation. were kept in the dark at a temperature of either 23 or 28’5°C in Danieau’s solution at a confluency of 50 embryos per plate.

Embryos older than 24 hpf were maintained in embryo medium (Danieau’s solution) containing 1% 1-phenyl-2-thiourea (PTU) (Sigma-Aldrich) to inhibit pigment formation. Embryos were staged as previously described (Kimmel et al., 1995).

For this study, we have used the following lines: *TgBAC(neurod1:eGFP)^nl1^* (Obholzer et al., 2008), *Tg(neurod1:kikume)* (generated in the lab, Bañón & Alsina, 2023), *Tg(brn3c:GFP)^s356t^* (Xiao et al., 2005).

### Microinjection

Long and very thin injecting needles were pulled in an electrophysiology puller (Sutter instruments model P-97) with the following protocol: *P*=200; HEAT=566; PULL=90; VEL=70; TIME=80. The tip of the needle was bevel broken using forceps and the needle was loaded with injection solution. Embryos were injected with 1 or 2 nl into the cell in one-cell-stage embryos using an Eppendorf Femtojet 4i microinjector.

For CRISPR Cas9 KO experiments, a mix containing 1.33 µL of each duplexed gRNA (3 dgRNA [crRNA+tracrRNA] in total, stock concentration 25µM), 0.5 µL (stock concentration 62 µM) of Alt R S.p.HiFi Cas 9 Nuclease v3 (#1081060) and 0.5 µL of nuclease-free water. Cxcl14 crRNAs were designed with the CHOPCHOP platform (https://chopchop.cbu.uib.no/) following published guidelines (Hoshijima et al., 2019; Wu et al., 2018). crRNA sequences targeting *cxcl14* for CRISPR-Cas9 KO: CATTTATTCGCTCAACACAG, GCACTAGGAACCTTGTCAAG and GGGGATGAATCGCTGTAGTA. Scrambled crRNAs (randomization of antibiotic resistance sequences) for CRISPR-Cas9 experiments: GGCTCCTTGCTCGCGACTTG, GGCGACCGTGGCCGGAACGA and GCCTGCGCGTACCGACTCGG. All crRNAs were ordered from Integrated DNA Technologies (IDT). Each crRNA was mixed with Alt-R™ tracRNA (1:1, IDT) to obtain duplexed gRNA.

For Tol2 injections (hsp70I:cxcl14-IRES-CAAX-GFP) a mix of 1 µL of Tol2 RNA at 175-200 ng/µL, 3 µL of plasmid at 50 ng/µL and 6 µL of dH_2_0 was prepared. 1 nL of this mix was injected into the cell of one-cell-stage embryos.

### Genotyping *cxcl14* F_0_ KO (crispants)

Genomic DNA extraction of embryos was performed using the N-Amp extraction kit (XNAT2 Extract-N-Amp Tissue PCR Kit XNAT2-1KT). PCR protocol was as follows: Fw primer: ACAGAGTTTATCATCAAACCG, Rv primer: ACTTTAACTTCGCAAATATGCC. An initial denaturation at 95°C 3 min; 35 cycles of 94°C 30 s; 58°C 30 s; 72°C 30 s; 72°C 7 min; hold at 4°C. Heteroduplex mobility assay was performed (electrophoresis was carried out on a 3% agarose gel, at 120V for 1h 30min), followed by Sanger sequencing of the samples to confirm mutations.

### Photoconversion

Photoconvertible *Tg(neurod:kikume)* embryos were maintained in the dark to avoid spontaneous photoconversion. Circular ROIs of the desired size were drawn using the Zeiss ZEN 3.11. Bleaching option of the software was employed for photoconversion. UV 405 laser power set to 10% and conducted in 3-4 iterations at a speed of 200 Hz. An image prior and post photoconversion was obtained enabling the confirmation of a correct photoconversion.

### Photoablation

An SP5 inverted Leica Multiphoton confocal microscope was used with Mai Tai multiphoton activated [Mai Tai BB DeepSee (Spectra Physics) tunable (710-990 nm) pulsed laser], humidity 4%, temperature 20°C. A BS/RLD mirror and SP715 filter were used with a 910 nm laser at 42% power, at 20× zoom with an HC PL APO 20×/0.75 immersion objective (506191, Leica) because no ROIs can be used in this confocal in multiphoton mode. PMT was employed and the pinhole kept completely open (600 nm). Scans ranged from three to six until a bubble formed (indicating destroyed tissue). Scan speed was 200-400 Hz, frame size 1024×512, bidirectional scanning was on, with line and frame average 1.

### 4-Di-2-ASP staining and injection into the otic vesicle

Embryos were maintained as described until 72 hpf, anesthetized using MS-222 (Tricaine) (42 µL of Tricaine at a stock concentration of 4 mg/mL per 1 mL of embryo medium) and placed in a lateral position on top of a 3% agarose-coated petri dish. Using the same setup as the one described in the microinjection section, 5 nL (20 ng/mL) of 4-Di-2-Asp (Sigma-Aldrich, D3418) were injected into one of the otic vesicles of the embryos. To label hair cells from neuromasts embryos were then incubated in a 1:1000 dilution of 4-Di-2-Asp (1 µL of 20 mg/mL in 1 mL of Danieau’s solution) shaking for 40 minutes. After 3 washes with Danieau’s solution, the embryos were returned to the incubator until the imaging step.

### *In situ* hybridization of whole-mount embryos

Embryo whole-mount *in situ* hybridization protocol was adapted from (Thisse and Thisse, 2008). Antisense riboprobes were generated by *in vitro* transcription using DIG or FLUO-dNTPs of cloned cDNA or directly from PCR-amplified fragments, using primers containing the promoter sequence of T7 or SP6 sequences at the 5’ ends of the reverse primer. Fixed embryos were dehydrated with methanol and stored at -20°C. After rehydration, permeabilization with proteinase K (diluted 1:1000 from a stock of 10 mg/mL, Invitrogen), embryos were incubated with the hybridization buffer (50% FAD, 25% SSC, 250 mg torula RNA, 1,25 mg heparin, Tween 0.1%) at 70°C for 2h. Incubation with the riboprobes diluted in hybridization buffer at a concentration of 0.5 ng/µL (DIG-labeled probes) or 2ng/µL (FLUO-labeled) was done overnight at 70°C. Embryos were washed and incubated with blocking solution (2% blocking reagent (Roche, 11096176001) 10% neutralized goat serum in 1x MABT) for 1h. Incubation with anti-FLUO-POD (1:400), anti-DIG-POD (1:1000), anti-FLUO-AP (1:2000) or anti-DIG-AP (1:2000), respectively, was performed O/N at 4°C. Fluorogenic *in situ* hybridization probes were developed with TSA Fluorescein and Cy3 (Akoya, NEL753001KT), respectively, for 1h. Chromogenic *in situ* hybridization probes were developed with NBT/BCIP. Signal development was assessed visually, time ranging in a probe-dependent fashion. After stopping the reaction with dH_2_O, embryos were kept in PBST/Glycerol 1:1 and flat-mounted for imaging.

### Embryo immunostaining

Embryos were fixed at the desired stage in 4% formaldehyde, washed 3×5 min with PBST 0.1% and permeabilized with proteinase K (diluted 1:1000 from a stock of 10 mg/ml, Invitrogen). Larvae were washed 2×10 min in PBT 0’1%, post-fixed with 4% PFA, and blocked in 10% neutralized goat serum and 2% bovine serum albumin in PBST for 2 h at room temperature. Embryos were incubated overnight at 4°C with antibodies diluted in blocking solution. Primary antibodies used: rabbit anti-GFP (1:400; Torrey Pines, TP401), mouse anti-HCS-1 (1:25, Developmental Studies Hybridoma Bank, AB_10804296). After washing with PBST, embryos were incubated with secondary antibodies conjugated with Alexa Fluor^®^ 488, 594 or 633 (1:500 dilution in blocking solution; Invitrogen, A-11029, A-21206, A-11037, A-21070 or A-21053).

### Confocal imaging

Live zebrafish embryos were anesthetized with MS-222(Tricaine) at the desired stage. Fixed or live zebrafish embryos were mounted 45° from completely lateral or dorsally in 0.8% low melting point agarose (Ecogen, 90002155) on a 35 mm glass-bottom Petri dish (MatTek, P35G-1.5-14-C). Images and time series were acquired with the LSM980 inverted confocal (Zeiss). Each channel was acquired rastering by line, bidirectionally, using the 40x immersion objective (Zeiss, 420862-9970-799, NA 1.2) with silicone oil. A z-step of 1 µm was selected. The image format was 1024×1024 or 2048×1024 and the scan speed ranged between 1.89 to 3.77 sec / frame. 488 nm and 561 nm laser excitation varied from 1 to 10% for *in vivo* acquisitions and up to 30% for fixed samples. The same settings, laser power and gain were maintained among embryos of the same experimental unit. Multi-position experiments (timeseries with multiple embryos) included up to 10 embryos separated by a time frame no longer than 20 mins. Airyscan function was used for imaging hair cells labelled with 4-Di-2-ASP and Otoferlin antibody to obtain higher spatial resolution images.

### Image processing

FIJI software was used to analyze confocal images. Image brightness and contrast were adjusted linearly. MAX projections of the whole z-stack or specific slices were done. Gaussian blur, subtract background (50 pixels radius) and/or minimum filter were applied to images for visualization purposes only. Data quantification was obtained from raw images. When comparing signal intensity in different conditions (i.e. 4-Di-2-ASP in control and crispant embryos), the same values of linear modifications were applied to all images. 3D drift correction (Fiji plugin) was applied to timeseries acquisitions, to correct tissue displacement at a pace of 4-10 µm/h.

### 3D reconstruction and quantification of axons

FIJI Single Neurite Tracing (SNT) from Neuroanatomy (https://imagej.net/plugins/snt/index) was used to trace axonal arborizations of z-stack images from 48 hpf embryos. ROIs containing an axon bundle directed to each crista were defined, starting from the most medial slice where the bundle started from somas to the tip of the most lateral branch. The bundle was analysed as a single axon. The bundle was traced up to the initiation of branches, and each branch was traced individually. Quantifications were performed using the analysis tools of SNT (cell-based analysis).

### Otoferlin quantification

From each crista analyzed (Fig. 7), lines were hand-drawn using the Fiji line tool, from the most apical tip of a hair cell to the base using the channel for HC membranes (*Tg(brn3c:GFP)^s356t^)*. 6 random HCs were selected from each crista. The intensity of Otoferlin signal was then plotted using the plot profile function in FiJI and the mean intensity of apical and basal regions were calculated using Excel. Comparisons and plots of the apical/basal ratio were performed using RStudio.

### Behavioral analysis

Fish were injected as previously described with either *cxcl14* or scrambled (control) gRNAs. Sibling larvae of both were kept at 28.5°C until 5dpf. Individual larvae were placed in wells of a 96-square-well plate containing 200µL of embryo water to be acclimatized for 2h before the behavior analysis (24 cxcl14 crispant and 24 control larvae per plate). Behavior analysis was performed using the DanioVision Observation Chamber (Noldus Information Technology, Wageningen, Netherlands) linked with the EthoVision XT17 software. A steady flow of water was supplied to the chamber and maintained the larvae at 28.0°C during the whole recording. Camera resolution was set at 1280 × 960 and the frame rate was set at 25 frames per second. The experiment underwent with the light on with an intensity of 5%. The experiment consisted in a startle response test elicited by 10 mechanical taps at 90-second intervals. The taps were generated using an inbuilt solenoid positioned at the base of the DanioVision chamber, with the stimulus intensity set to the highest level (level 8).

EthoVision XT17 analyzes changes in pixels per frame. Activity within arena was quantified as the percentage of pixels moving in each well. Distance moved was quantified as the total movement of the center point detected for each larva. From raw individual tracking data, the startle response strength was calculated as the distance moved within the second post-stimulus, normalized by the baseline distance move per second during the five seconds before the stimulus (the baseline activity was subtracted to the distance moved during the startle stimulus). Distance travelled per second was quantified by summing frame-to-frame displacements in one second (sample rate = 1 image every 0.04 seconds). Each individual was measured for all 10 taps, then normalized and averaged and, finally, the group means and standard deviation were calculated.

### Statistical analysis and plots

Graphs were generated with Graphpad Prism 9 software or RStudio (version 2024.12.1). Shapiro-Wilk normality tests were performed to categorise the distribution of data into parametric and non-parametric tests. For not normally distributed data we used the Mann-Whitney-Wilcoxon test. In the analysis of startle response strength of the behavioral test, a linear mixed effect analysis was used to test for effects from the different stimuli repeats. Values are expressed as mean ± standard deviation.

### Single cell RNA sequencing

The scRNA-seq datasets have been previously published (Olson et al., 2024; Woodruff et al., 2025) and deposited in the Gene Expression Omnibus (GEO) database under accession codes: <u>GSE266312</u>, <u>GSE264323</u>, and <u>GSE24072163</u>. The LL cluster was identified by expression of known lateral line markers (*hoxb5b*, *gfra1b*, and *hmx4*) but lack of marker expression of other cranial sensory neurons (*irx1a* and *phox2bb*). This cluster was then subjected to unsupervised subclustering, which yielded 6 distinct subpopulations. To identify genes enriched in each subpopulations, differential expression (DE) analysis was performed with a minimum difference threshold set at 20% and logfc threshold = 0.2 using the ‘FindAllMarkers’ function of Seurat. We then used top 10 DE markers to find the identity of each subcluster.

R-code used in this study is available on GitHub: https://github.com/anechipor/Rumbo_et_al_2026

## Acknowledgments

We thank members of the laboratory for insights and critical discussions (Francesca Manocchio, Clara Gordillo, Gonzalo Ortiz, Carolyn Engel-Pizcueta, Lydvina Meister). CRG-ALMU microscopy facility staff for technical support in image acquisition in Leica SP8 and Zeiss LSM980 systems. We would also want to thank Dr. Manuel Irimia and Cristina Rodríguez-Marín for letting us use the DanioVision Observation Chamber and the help provided.

## Author Contributions

Conceptualization of the project: M.R, A.B, B.A; Most experiments were done by M.R and A.B. A.V.N performed the scRNA-seq analysis; Manuscript writing was done by M.R and B.A. Manuscript revisions were done by A.B, M.R, A.V.N, B.A.

## Funding

This work was supported by Ministerio de Ciencia e Innovación AEI-BFU2017-82723P, PID2020-117662GB-I00 (FEDER) to B.A. and the Unidad de Excelencia María de Maeztu, Agencia Estatal de Investigación (CEX2018-000792-M). M.R. is recipient of the PRE2021-097774 predoctoral fellowship and A.B. was a recipient of the predoctoral fellowship ‘Formación de Profesorado Universitario (FPU)’ from the Spanish Ministerio de Universidades (FPU17/03287). A.V.N. is supported by an award from NIH: NS143805 (NINDS).

## Notes

### Competing Interest Statement

The authors have declared no competing interest.

