## Supplementary Figures for "The Cxcl14 chemokine defines pioneer axon guidance and early circuit assembly in the inner ear"

### Supplementary Material and Figure legends

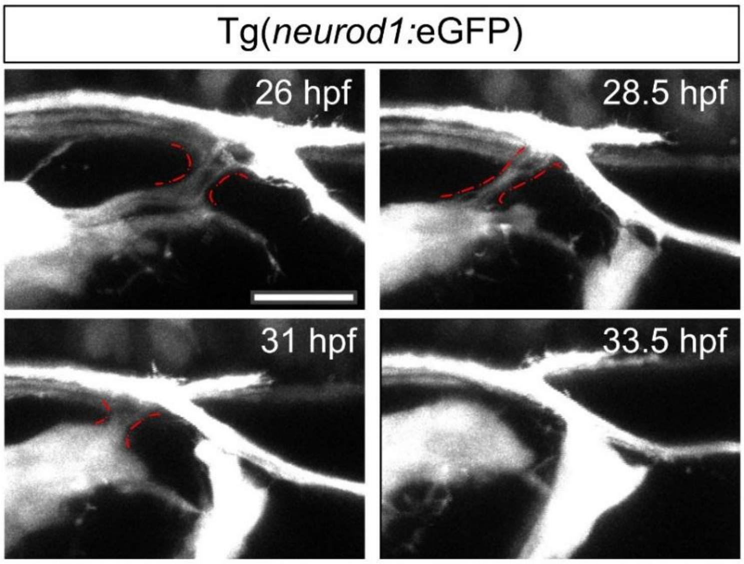


**Suppl Fig. 1. Disassembly of SAG axons and pLL contacts.** Representative confocal stills of a timeseries from 26 to 33.5 hpf of TgBAC(*neurod1:eGFP*)$^{nl1}$. Axonal contacts are marked with red dotted lines. Scale bars = 20 µm.

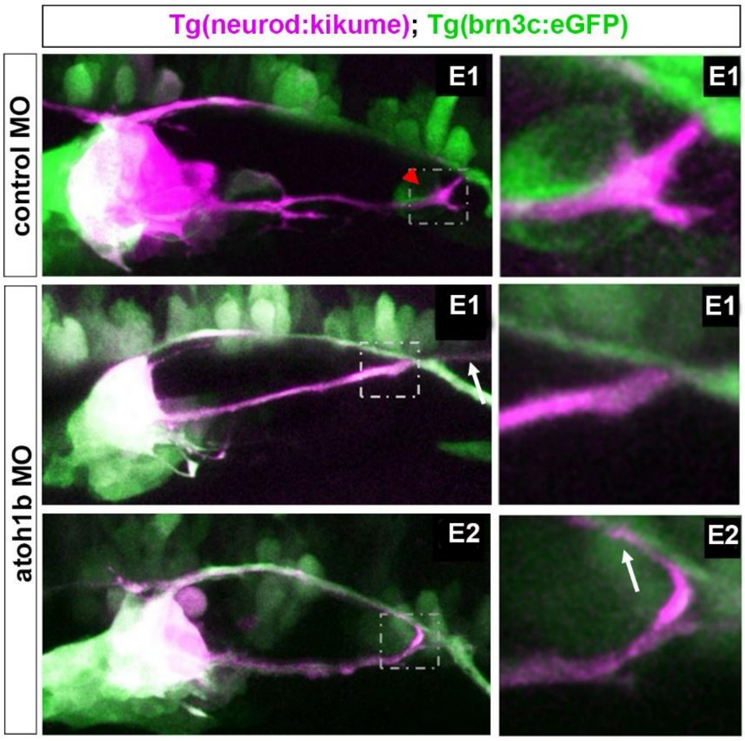


**Suppl Fig.2. atoh1b depletion results in loss of HC contact by SAG pioneer axons and a consequent extended growth.** Representative confocal images of control and *atoh1b* morphants. SAG pioneers are labelled through Tg(*neurod1*:*kikume*) and photoconverted (magenta) and HCs are labelled in by Tg(*brn3c*:*eGFP*)$^{s356t}$ in green. Red arrowhead depicts axon swelling. White arrows depict the extended growth of pioneer axons in atoh1b morphants.

**Video 1.** Growth of pioneer axon to the posterior region of the OV from 20 to 23 hpf. Red arrowheads label axon growth tips. Scale bar = 20 µm.

**Video 2.** Branching of pioneer axons from 22 to 24.5 hpf. Red arrowheads indicate branches. Scale bar = 20 µm.

**Video 3.** Crawling of neuroblasts along the pioneer axons to form the posterior lobe of the SAG from 30 to 36 hpf. Scale bar = 10 µm.

**Video 4.** Example of cxcl14 crispant embryo (E1 in Fig. 4b) from 22 to 25 hpf. Red arrowheads indicate SAG pioneer axons. Blue arrowheads indicate axons of the pLL entering into otic territory. Scale bar = 20 µm.

**Video 5.** Example of cxcl14 crispant embryo (E2 in Fig. 4b) from 22 to 25 hpf. Red arrowheads indicate SAG pioneer axons. Scale bar = 10 µm.

**Video suppl 1.** Disassembly of SAG-pLL axon contacts from 26 to 33.5 hpf. Scale bar = 10 µm.
